# The *Cancermuts* software package for the prioritization of missense cancer variants: a case study of AMBRA1 in melanoma

**DOI:** 10.1101/2022.05.23.493014

**Authors:** Matteo Tiberti, Luca Di Leo, Mette Vixø Vistesen, Rikke Kuhre, Francesco Cecconi, Daniela De Zio, Elena Papaleo

## Abstract

Cancer genomics and cancer mutation databases have made a wealth of information about missense mutations found in cancer patient samples. Contextualizing by means of annotation and predicting the effect of amino acid change help identify which ones are more likely to have a pathogenic impact. Those can be validated by means of experimental approaches that assess the impact of protein mutations on the cellular functions or their tumorigenic potential. Here, we propose the integrative bioinformatic approach *Cancermuts*, implemented as a Python package. *Cancermuts* is able to gather known missense cancer mutations from databases such as cBioPortal and COSMIC, and annotate them with the pathogenicity score REVEL as well as information on their source. It is also able to add annotations about the protein context these mutations are found in, such as post-translational modification sites, structured/ustructured regions, presence of short linear motifs and more. We applied *Cancermuts* to the intrinsically disordered protein AMBRA1, a key regulator of many cellular processes tightly (de)regulated in cancer. By these means, we classified mutations of AMBRA1 in melanoma, where AMBRA1 is highly mutated and displays a tumor-suppressive role. Next, based on REVEL score, position along the sequence and their local context, we applied cellular and molecular approaches to validate the predicted pathogenicity of a subset of mutations in an *in vitro* melanoma model. By doing so, we have identified two AMBRA1 mutations which show enhanced tumorigenic potential and are worth further investigation, highlighting the usefulness of the tool. *Cancermuts* can be used on any protein targets starting from minimal information, and it is available at https://www.github.com/ELELAB/cancermuts as free software.

## Introduction

Recent advances in cancer genomics have been leading to increased information on cancer mutations. Resources as Genomic Data Commons (GDC) [1] store information from different studies from cancer genomic initiatives, such as The Cancer Genome Atlas [2] and the Therapeutically Applicable Research To Generate Effective Treatments (TARGET) initiative (https://ocg.cancer.gov/programs/target). Databases such as the Catalogue of Somatic Mutations in Cancer (COSMIC) [3] or cBioPortal [4, 5] are also a useful resource to mine cancer mutations. Providing annotations and predictions, which may help discriminate between mutations with or without a pathogenic impact, is still an open challenge [6-9] and a field in need of urgent investigation. This could be assessed by experimental approaches to determine the impact on protein cellular functions or the tumorigenic potential deriving from the alteration. It is also noteworthy that genomic-related changes in coding regions leading to amino acidic substitution(s) can possibly result in alterations of the protein product in terms of stability, key post-translational modifications regulating protein function, or even interactions with other proteins. Structure-based methods can be applied to assess these different functional layers [10-16] even though they have a limited scope, especially if the target protein includes large intrinsically-disordered regions or regions enriched in low-complexity sequence patterns. Furthermore, the context in which any mutation is found is also relevant, as it can be indicative of putative effects of a mutation. For instance, it can be useful knowing whether a certain substitution falls within a binding site for another protein, or whether it is located within a structured region, or whether the substitution can abolish or even introduce a new post-translational modification.

To give an easily accessible overview of (i) the distribution of mutations in a gene, (ii) the pathogenicity scores and (iii) the annotations along the protein sequence, we have created *Cancermuts*, a Python package that streamlines the collection and annotations of cancer mutations located in the coding region of a gene of interest, e.g. mutations that will impact its protein product. The information is superimposed with different layers to help make informed decisions on which mutations are more likely to be functionally damaging and worth further investigation by either computational or experimental approaches.

To validate *Cancermuts* in a cancer study, we focused on the tumor suppressor gene *AMBRA1* (autophagy and beclin 1 regulator 1). Initially discovered to be involved in correct embryogenesis, especially during brain development, in mouse congenital malformations as well as in human neurological disorders [17, 18], AMBRA1 is mostly known for its role in autophagy activation [17, 19]. New cancer-related roles for AMBRA1 have emerged over the years, particularly with regard to cell proliferation and tumorigenic potential [20]. More recently, the role of AMBRA1 as tumor suppressor has been further extended, as by its regulation of cell cycle by Cyclin D1 stabilization (*via* interaction with the E3 Ubiquitin ligase DDB1-Cullin4) [21-23] and of malignant invasiveness (through focal adhesion kinase FAK1 hyperactivation) [24]. Such a vast range of functions is intertwined with the ability of AMBRA1 to interact with molecular partners [17, 19]-[23, 25]-[33] and undergo post-translational modifications (PTMs) [19, 34, 35], and deeply relies on its intrinsically disordered structure.

In this study, we used *Cancermuts* on AMBRA1, allowing to identify putative cancer mutations of interest to be further validated experimentally. The prediction of pathogenic mutations of AMBRA1 and their *in vitro* validation have been carried out in melanoma, the most aggressive and lethal form of skin cancer, in which AMBRA1 not only bears an anti-tumorigenic function [24], but also displays high mutation rate [21].

## Results

### Design and implementation

*Cancermuts* is designed as a Python package with an easy-to-use and well-documented programming interface (API) (**Fig. 1**). It is suited to researchers with basic programming Python skills and can be used, for instance, in popular interactive Python interfaces, such as Jupyter notebooks or Google Colab, as well as in standalone Python scripts, or integrated in more complex workflows.

**Figure 1.**
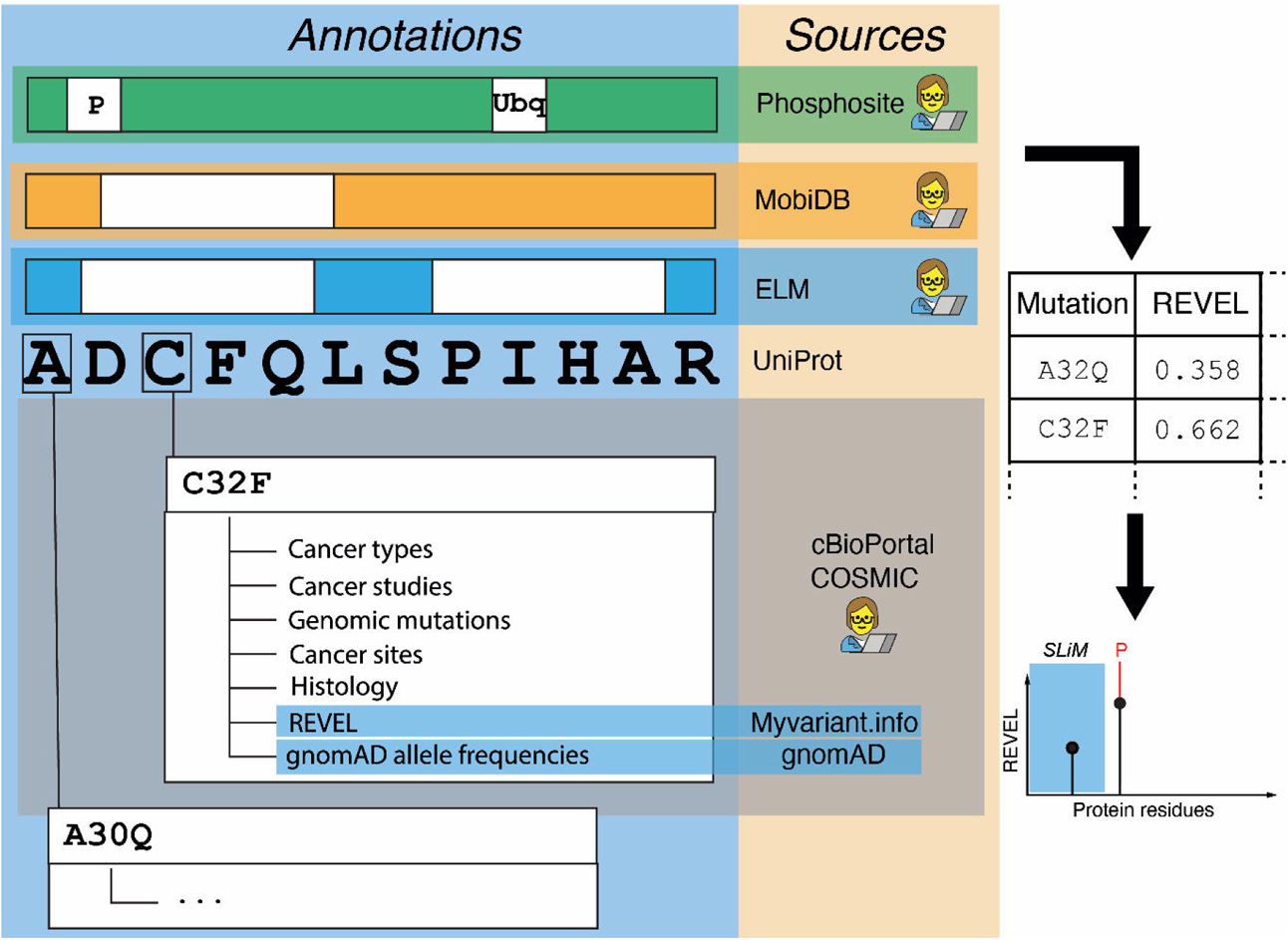
Schematic representation of the Cancermuts workflow. The figure shows on the left the different layers of evidence that Cancermuts supports. The Uniprot main isoform sequence is the basis on which either per-position or per-sequence annotations are performed (post-translational modifications, structure definition, linear motifs) that can be provided by one of the sources and by manual user annotation. The sequence can also be annotated by downloading cancer-related mutations and relative metadata, including REVEL scores and gnomAD allele frequencies. Mutations can be supplied by manual annotation as well. Once the information has been collected, it can be summarized as a table (right) and as a plot (bottom right).

*Cancermuts* only requires basic information about the gene of interest, namely either its IDs or its protein product IDs, such as Uniprot accession ID [36]. Using the *Cancermuts* API, the user is expected to download the protein sequence first by providing relevant database IDs (**Fig. 1**). This can then be annotated with protein missense mutations from cancer mutation or genomics databases, as well as with further annotations regarding the protein sequence itself and the identified mutations (see below for details). Both mutations, e.g. from patient-cohort studies, and annotations can also be provided from custom user-designed input files, allowing integration in the annotation pipeline. The tool has been designed to annotate somatic mutations and especially focuses on single nucleotide variants. Once the data collection has been performed, *Cancermuts* provides the researcher with textual and graphical representation of the mutations to explore the data and help with their interpretation. All the obtained data can be converted to a simple Pandas data frame, which can then be manipulated as desired, including saving it as a table (CSV) file. *Cancermuts* also includes facilities to represent the annotation as a publication-ready stem plot which can be thoroughly customized.

*Cancermuts* interacts with different freely available resources for sequence-based annotations as detailed below. The package is designed to be modular and easily extendable, should other annotations be of interest in the future.

The current release interacts with the cancer databases COSMIC [3] and cBioPortal [4, 5] to retrieve cancer-associated mutations, allowing on-the-fly or local access, respectively, along with data filtering starting from minimal information about the gene of interest. It is also possible to filter for cancer type or study. Some of the metadata from these databases are kept as annotations.

In addition, *Cancermuts* retrieves a score (ranging between 0 and 1) for pathogenicity of the mutations based on 13 individual predictors that have been combined using a random forest approach within the REVEL framework [37]. The user can deduce the threshold value to associate with pathogenic mutations based on specific case studies and benchmarking. We recommend applying a cut-off of 0.40 in case additional information are lacking, i.e. the one that guarantees a good compromise between specificity (0.85) and sensitivity (0.81) [37].

The tool is also able to query gnomAD [38], a collection of harmonized whole-genome and - exome sequencing data. gnomAD works as a proxy of the healthy population for allele frequency. This annotation can be used, for example, to discard some of the mutations from further analyses. Indeed, if a mutation occurs with high frequency in non-tumoral samples, it is unlikely to have a strong pathogenic impact.

*Cancermuts* allows to store annotations for functional short-linear motifs (SLiMs) that might be related to protein regulation or interaction. This is done interacting with the ELM database [39] or providing input annotations from other sources. Information on putative PTMs are provided by querying PhosphoSitePlus [40] and can additionally be provided by the user in case additional annotations not covered in the database (but experimentally proven) are available.

*Cancermuts* also allows to annotate the propensity to disorder or structure using MobiDB. Additional custom annotations regarding structure propensity can be provided by the user through a specific formatted CSV file (see GitHub repository and user guide).

*Cancermuts* has been designed to be applicable to any protein product and does not require structural information. It is especially interesting for intrinsically disordered proteins or domains, along with low complexity repeats for which structure-based methods currently available to predict the effect of mutations are not easily transferable or challenging to apply. Structural annotations can be facultatively added, whether available.

### Case study: AMBRA1 mutations in melanoma

AMBRA1 is a large, mostly disordered protein, with a canonical UniProt sequence of 1298 residues. The intrinsically disordered nature of the protein, along with its high plasticity, probably due to its several protein-protein interactions and post-translational modifications (PTMs) [17-35], make of AMBRA1 a good candidate to link together different intracellular processes. Notably, many types of cancer, including malignant melanoma, where AMBRA1 has been shown to play an anti-tumorigenic role [24], show genetic alterations in *AMBRA1* [21-23]. Indeed, AMBRA1 – when compared to other cancer studies, shows the highest mutation rate in skin cancers, such as melanoma [21].

Due to its structure, propensity to interact with other proteins and cancer-related functions [21-24], we sought to apply Cancermuts to *AMBRA1* in order to predict the pathogenicity of its mutations in melanoma.

### Cancermuts *for AMBRA1:* in silico *analysis*

We have downloaded all available mutations for AMBRA1 for melanoma-associated mutations from COSMIC and mutations associated with melanoma studies from cBioPortal on April 8^th^, 2020. We have annotated this information with all the available annotations in *Cancermuts* as well as integrated them with manual annotations. These include information about SLiM binding sites and PTMs known in literature but unavailable on the databases on the date the pipeline was run, as well as more about predicted structural regions of AMBRA1 (see GitHub repository). By using a model based on AlphaFold2, we have predicted residues ∼7-203 and 857-1040 (381 residues: ∼29% of total sequence length) of AMBRA1 to be structured regions, including a region with a β-propeller fold (**Fig. 2A**). We have saved all the collected information in a CSV table (see **Table S1**) and provided a graphical support (**Fig. 2B**).

**Figure 2.**
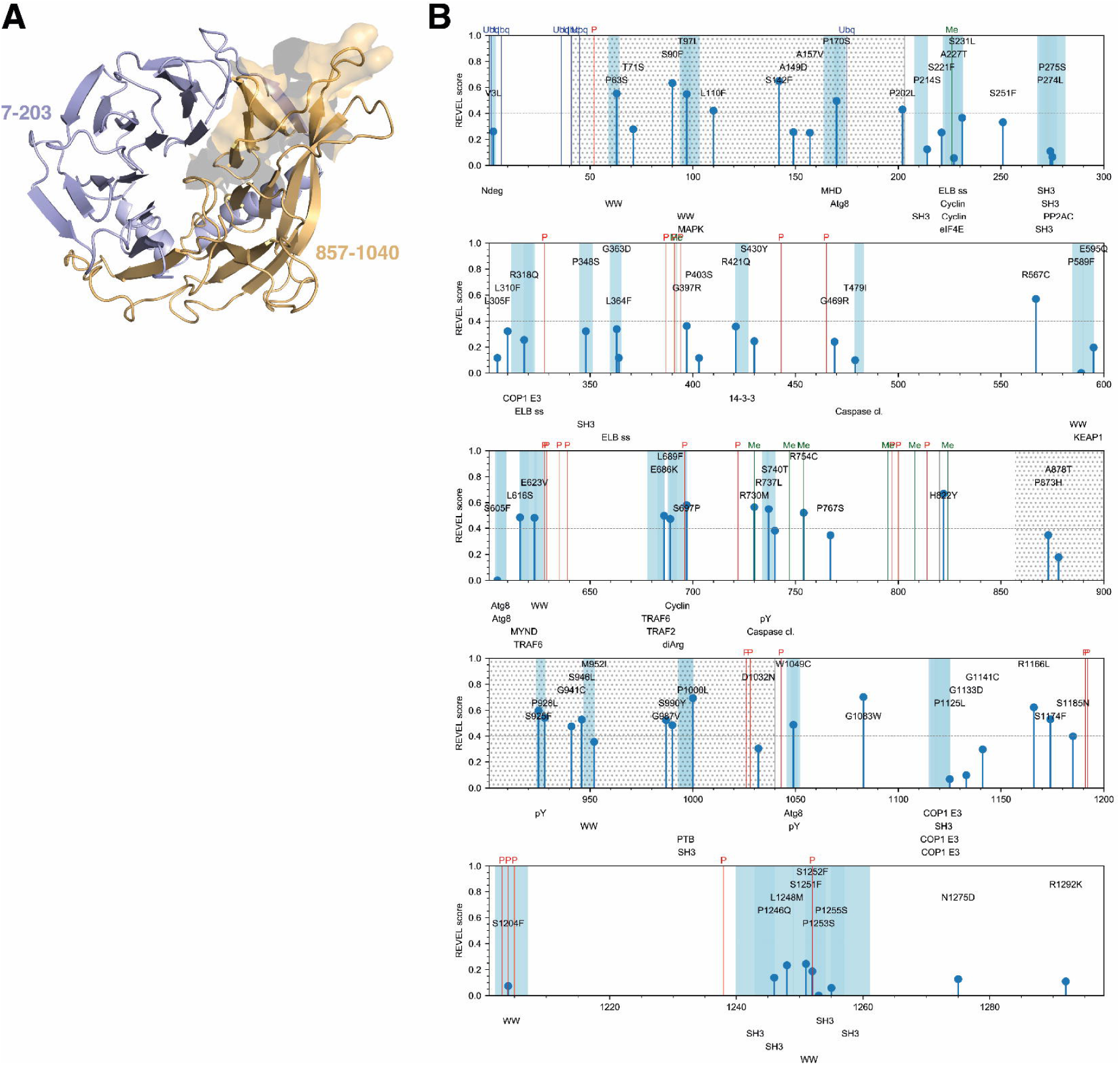
Predicted structured regions of AMBRA1 and identified cancer mutations. (**A**) predicted structured regions of AMBRA1 including the β-propeller domain. The two residue stretches the structure is composed of are shown in blue (7-203) and orange (857-1040), respectively. The part of this structured region against which the AMBRA1 antibody we have used has been raised against is also highlighted by showing its molecular surface (top right). (**B**) Melanoma-related mutations and corresponding annotations as collected by Cancermuts. Plots follow the main Uniprot sequence numbering; each mutation is annotated as a stem the height of which is proportional to the identified REVEL score (with 0 for those that could not be annotated). Blue shades and corresponding bottom labels refer to linear motifs as annotated by ELM. Post-translational modifications are shown as colored vertical lines. A grey dotted pattern was used to represent the predicted structured β-propeller domain of AMBRA1. Predicted SLIMs that do not overlap with mutations have been hidden from this plot for ease of visualization.

Overall, our analysis identified a total of 73 melanoma-associated non-neutral mutations (**Table S2, Fig. 2B**), 70 of which deriving from single-nucleotide substitutions and 3 from multiple nucleotide substitutions (P589F, S605F, P1253S). As the REVEL score is only available for single substitution, these could not be assigned any score. Of such mutations, 40% (28) displayed a REVEL score below significant threshold (**Table S2, Fig. 2B**). Out of the identified protein mutations, the genomic alterations associated to 54 (∼74%) were found compatible with UV-induced DNA damage (**Table S3**). Overall, about 39% of the identified mutations were found to be in putative structured regions, displaying no general preference for such regions to accumulate mutations. Nonetheless, out of the 28 mutations predicted as damaging for REVEL, the majority (15) was found within the predicted structure regions, whereas only 13 were found within the disordered parts of the protein, which covers nearly 60% of the sequence. Therefore, at least for this specific case, a damaging mutation is more likely to be found in a predicted structured region. Pro and Ser were by far the most mutated residue types (17 for each, respectively), followed by Arg (8), Gly (8), and Leu (7). Not surprisingly, the most frequent mutation in the dataset was Ser to Phe (10 occurrences) and Pro to Ser (9), followed by Pro to Leu (5) and Leu to Phe (5), with all the other substitutions being far less frequent (two occurrences or less). The mutation frequencies downloaded by gnomAD did not allow us to discard any of the identified mutations in this case.

We have annotated a total of 26 phosphorylation sites, 6 ubiquitylation sites and 7 methylation sites. Most of the PTMs sites are localized in unstructured regions of the protein, where they could be more accessible to kinases or other proteins responsible for their modification. Phosphorylations tend to be clustered in small groups, for instance in stretches 387-394, 628-639, 797-814, 1027-1043, 1192-1205, which may be important regulatory regions of the protein [19, 20, 26, 28, 29, 35]. Ubiquitylation sites are clustered in the first 50 residues of AMBRA1, while methylation sites are found mostly in the 730-824 region.

ELM identified several potential SLiMs to which interactors could bind close to mutation sites. It should be noted that SLiMs are defined in the context of disordered regions, while the SLiMs identified in *Cancermuts* are not filtered according to *Cancermuts*’ own definition of structured or unstructured regions, as such information could still be useful, depending on how trustable the disorder prediction or definition is, as well as considering order-disorder transitions in the protein structure. Interestingly, ELM also identified different TRAF6 ubiquitin ligase binding sites, the role of which in AMBRA1-mediated control of autophagy has already been described [19]. Other relevant binding sites include those for cyclins.

### Cancermuts *for AMBRA1:* in vitro *validation*

Among the identified mutations, several of the most pathogenic-predicted are included both in the N- and C-terminal predicted β-propeller regions (**Fig. 2B**). Based on the recent findings indicating that the N-terminus of AMBRA1 is involved in stabilization of AMBRA1 itself [25] and of Cyclin D1 [21-23], a result that we confirmed in BRAF^V600E^-mutated A375 melanoma cells silenced for *AMBRA1* by small interference RNA (siRNA; siAMBRA1 #1 and #2) (**Supplementary Fig. 1**), we characterized the *in vitro* effects of AMBRA1 mutations (REVEL score ≥0.4) within those mapping at the N-terminus of the protein (**Fig. 3A**). The list of these mutations, as well as interaction sites [19-23, 25-29] and PTMs [17, 25, 27, 30-33] of AMBRA1 are shown in **Fig. 3A**. Our analyses also include the A157V mutation which, bearing a REVEL score ≤0.4, and an amino acid substitution with a residue of similar type and steric incumbrance (A to V), is not predicted to be pathogenic. Re-expression of WT AMBRA1 has been used as reference. Our experimental settings consist of transfecting melanoma cells with a siRNA targeting the 5’-UTR region of *AMBRA1* (siAMBRA1#2) prior to mutant re-expression (**Fig. 3B**). Western blot (WB) analyses ruled out any possible effects of the mutated constructs on either the autophagy or apoptotic functions of AMBRA1, as respectively stated by lipidation of LC3 (LC3-II) (**Fig. 3C, D**), a *bona fine* marker for autophagy, and cleavage of the apoptotic marker CASP-3 (**Fig. 3C**). Instead, increased protein levels of Cyclin D1 were observed solely upon re-expression of the L110F mutant (**Fig. 3C, E**). Additionally, re-expression of L110F, and of P170S as well, resulted in hyperphosphorylation of SRC (a component of FAK1 signaling) at Y416 (pSRC-Y416), suggesting an active FAK1 signaling in both conditions (**Fig. 3C, F**). Interestingly, re-expression of both L110F (close to the DDB1-Cullin4 domain) and P170S (close to a predicted ubiquitination site) mutants resulted in poor detection of AMBRA1 constructs at protein level (**Fig. 3C, G**). On the other hand, no differences were depicted at mRNA level by RT-qPCR with respect to WT-expressing cells (**Fig. 4A**), hence suggesting possible effects on protein stability. Previously, AMBRA1/DDB1-Cullin4 interaction was shown to promote AMBRA1 stability by proteasome degradation [25]. To assess whether reduced AMBRA1 protein levels could result from protein degradation by either the proteasome or lysosome pathway, L110F- and P170S-expressing A375 cells were treated with proteasome (MG-132) (**Fig. 4B**) and lysosome (chloroquine, CQ) (**Fig. 4C**) inhibitors, respectively. However, in neither condition a rescue of protein levels was observed. The presence of protein aggregates was also assessed in insoluble fractions of mutant-expressing cells, however unsuccessfully (**Fig. 4D**). Surprisingly, when myc-tagged AMBRA1 constructs were re-expressed (**Fig. 4E**), and protein levels detected using an anti-myc antibody instead, the expression of the L110F and P170S mutants was comparable to WT-expressing cells, hence suggesting a failure of the anti-AMBRA1 antibody rather than effects of the mutants on protein stability (**Fig. 4E**). Levels of Cyclin D1 and pSRC-Y416 upon re-expression of myc-tagged mutants were also consistent with those observed in non-myc-tagged expressing cells (**Fig. 4E-G**).

**Figure 3.**
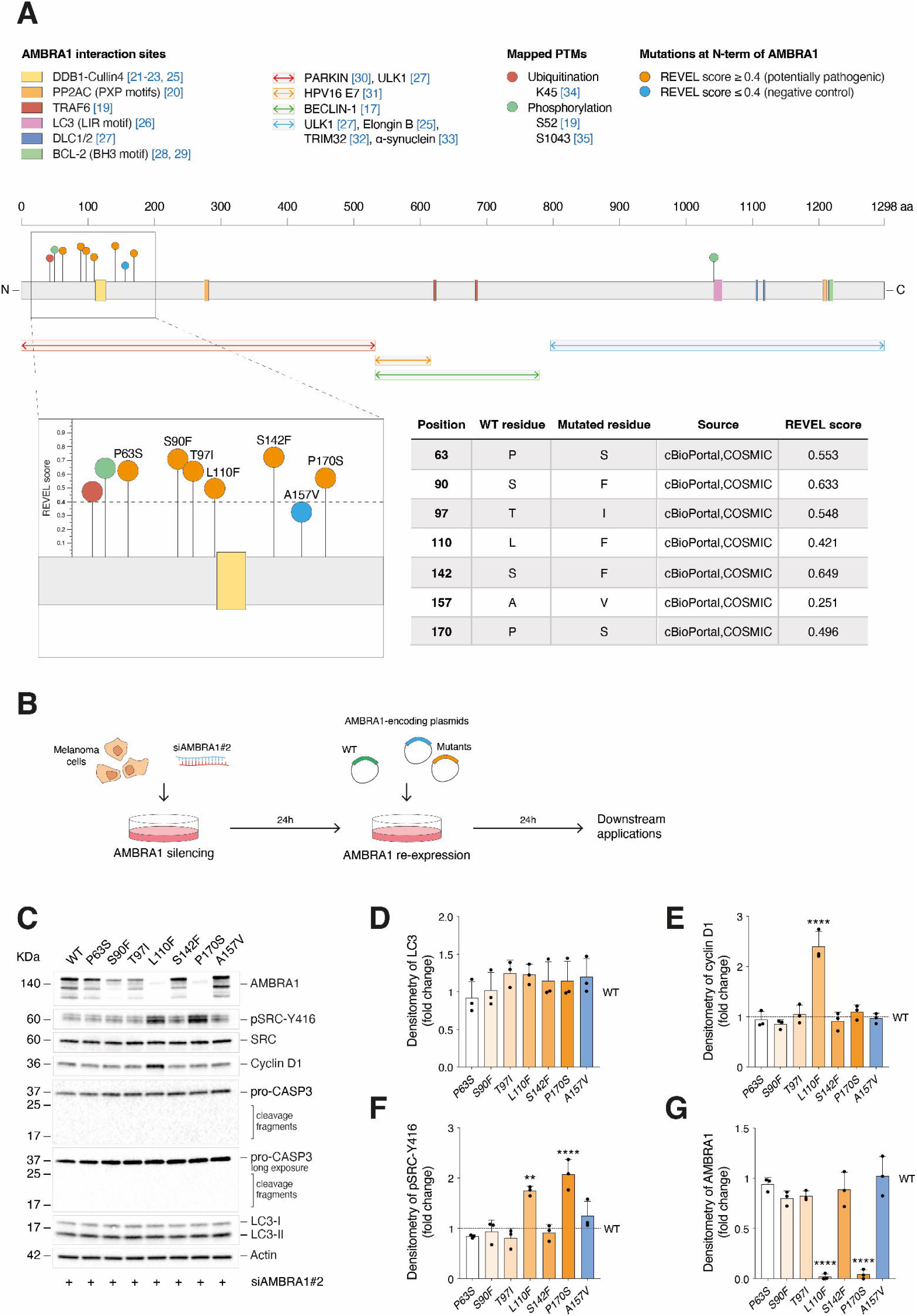
Analysis of the effect(s) of the point mutations of AMBRA1 on its functions. (**A**) Schematic representation of AMBRA1 including interaction sites and PTMs identified experimentally. AMBRA1 mutations are also mapped and specified in the table to the right. (**B**) Schematic representation of the transfection strategy. (**C**) WB analysis of pSRC-Y416, SRC, Cyclin D1, pro-CASP3 (including cleaved fragments) and LC3 (-I and -II) in A375 melanoma cells re-expressing the P63S, S90F, T97I, L110F, S142F and P170S AMBRA1 mutants. Re-expression of WT and A157V AMBRA1 was used as reference and negative control, respectively. AMBRA1 and Actin were detected as transfection and loading control, respectively. Images are representative of n=3 independent experiments. (**D-G**) WB quantifications are shown as fold change vs WT (represented by a dashed line) after normalization on the internal control Actin in (**D**) for LC3 (LC3-II/LC3-I ratio), (**E**) for Cyclin D1 (****p<0.0001 vs WT; one-way ANOVA), (**F**) pSRC-Y416 (pSCR-Y416/SRC ratio) (**p=0.0016; ****p<0.0001 vs WT; one-way ANOVA) and (**G**) for AMBRA1 (****p<0.0001 vs WT; one-way ANOVA). Data are expressed as mean ± SD.

**Figure 4.**
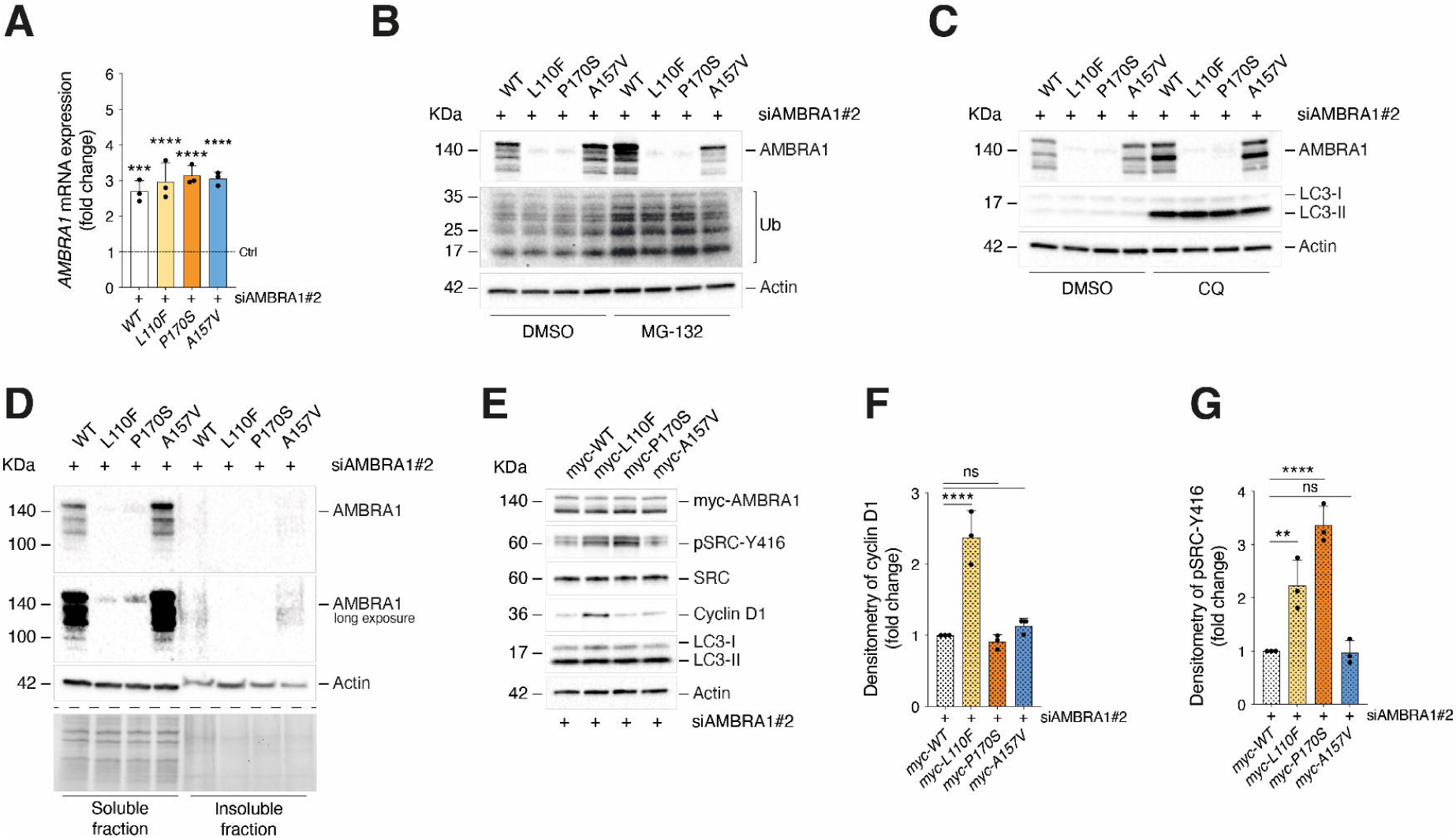
Analysis of the role of L110F and P170S mutants on AMBRA1 stability. (**A**) RT-qPCR analyses of *AMBRA1* upon WT, L110F, P170S and A157V re-expression. Data were normalized on *L34* and expressed as fold change vs non-transfected cells (Ctrl, indicated by a dashed line) ± SEM (n=3; ***p=0.0002; ****p<0.0001; one-way ANOVA). **(B)**24h after transfection of the constructs, A375 cells were treated with MG-132 (10 µM) or **(C)** CQ (40 µM) for 4h and WB analyses for AMBRA1 and Actin performed. Ubiquitylated proteins (Ub) and LC3-II accumulation were detected as positive controls for the treatments. Images are representative of n=4 independent experiments. (**D**) WB analyses of soluble and insoluble fractions upon mutant re-expression. AMBRA1 and Actin were detected (n=3). A representative gel activation is also shown. (**E**) A375 cells were re-expressed with AMBRA1-myc-tagged WT, L110F, P170S and A157V constructs and pSRC-Y416, SRC, Cyclin D1 and LC3 (-I and -II) detected by WB. AMBRA1 was detected using an anti-myc antibody as transfection control, whereas Actin as loading control. (**F-G**) Densitometry analyses for (**F**) Cyclin D1 and (**G**) pSRC-Y416 (pSRC-Y416/SRC ratio) were performed and normalized on Actin. Data are expressed as fold change vs myc-WT ± SD (n=3; ns=not significant; **p=0.0047; ****p<0.0001; one-way ANOVA).

The increased levels of Cyclin D1 and the hyperactivation of FAK1 signaling upon *Ambra1* depletion have been previously correlated to boosted proliferative rate and invasiveness of melanoma, respectively [24]. Although the increased Cyclin D1 levels observed upon L110F expression (**Fig. 3C, E**) did not correlate with changes in proliferation rate of A375 cells (**Fig. 5A, B**), the hyperphosphorylation of SRC (**Fig. 3C, F**) associated with a higher invasive capacity of A375 upon both L110F and P170S re-expression (**Fig. 5C, D**). No effects were observed upon re-expression of the negative mutant A157V. Such an effect was also unrelated to possible changes in cell viability (**Fig. 5E**). Consistently with previous data showing that loss of *Ambra1* promotes an epithelial-to-mesenchymal (EMT)-like phenotype in melanoma [24], re-expression of the L110F and P170S also improved mRNA expression of the EMT markers *Fibronectin* (*FN1*) (**Fig. 5F**) and *Vimentin* (*VIM*) (**Fig. 5G**).

**Figure 5.**
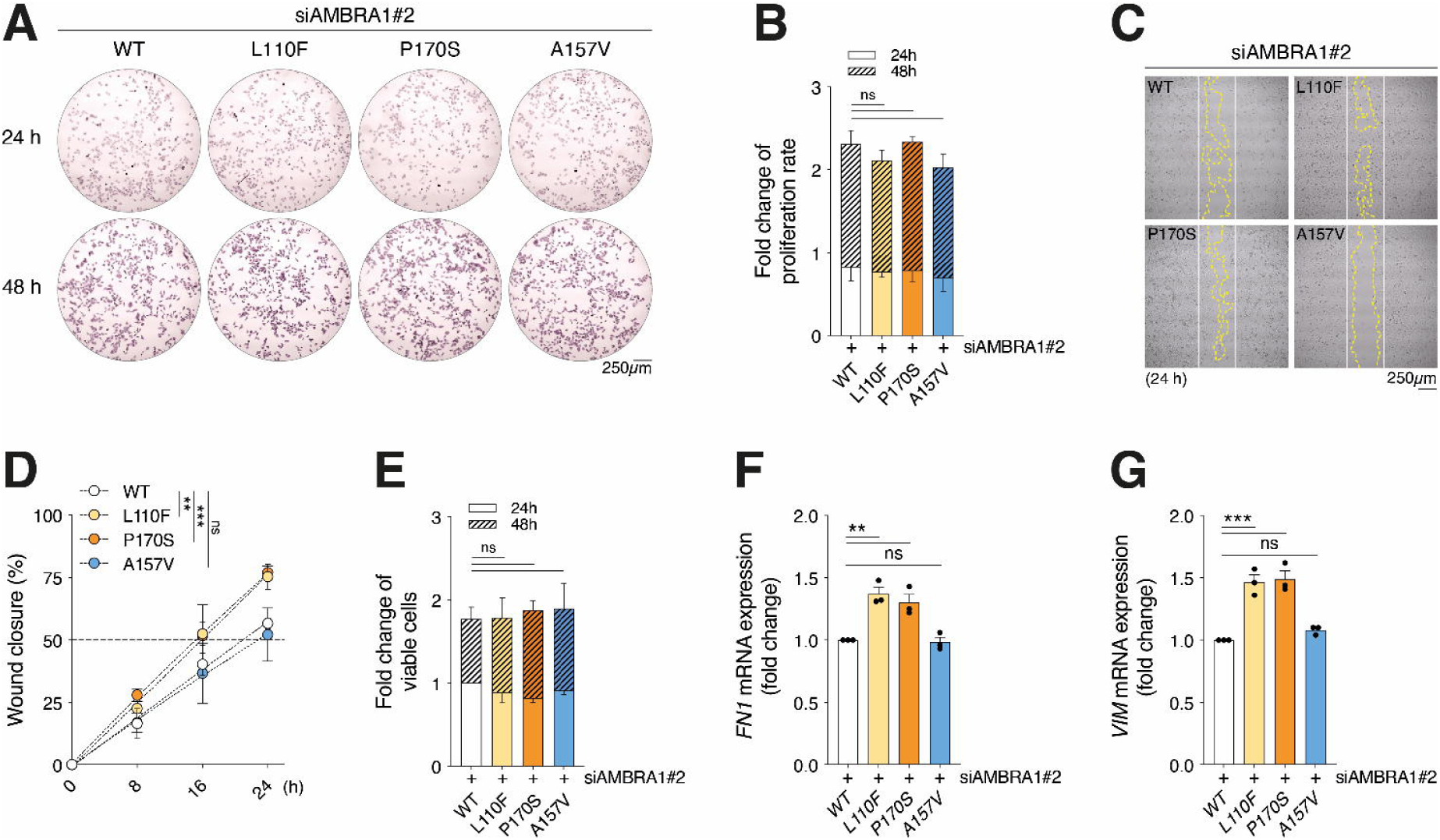
Analysis of the effect of L110F and P170S mutants on melanoma invasiveness. (**A**) Representative Crystal Violet staining of WT-, L110F-, P170S- and A157V-expressing A375 cells after 24 and 48h (n=3). (**B**) Quantification of the staining shown in (**A**). Data are shown as fold change ± SD with respect to control sample (WT at 24h) (n=3; ns=not significant; two-way ANOVA). (**C**) Representative images (n=3) of wound healing assay in mutant-expressing A375 cells at 24h. White and yellow lines outline wound edge at T_0_ and at the time indicated. (**D**) Quantification of wound closure is shown as percentage ± SD vs T_0_ at the times indicated (nC=C3; ns=not significant; **p=0.0018; ***p=0.0004; two-way ANOVA). (**E**) Cell viability of WT-, L110F-, P170S- and A157V-expressing A375 cells after 24 and 48h is expressed as fold change ± SD with respect to control sample (WT at 24h) (n=3; ns=not significant; two-way ANOVA). (**F**) RT-qPCR analyses of EMT markers *FN1* (n=3; ns=not significant; **p=0.0016 L110F vs WT; **p=0.006 P170S vs WT; one-way ANOVA) and (**G**) *VIM* (n=3; ns=not significant; ***p=0.0003 L110F vs WT; ***p=0.0002 P170S vs WT; one-way ANOVA) upon WT, L110F, P170S and A157V re-expression. Data are expressed as fold change vs WT ± SEM.

Comprehensively, our results indicate that, although differently, the expression of the AMBRA1 L110F and P170S mutants, predicted to be pathogenic, affects the invasive capacity of melanoma cells.

### Prediction of changes in folding free energy upon mutations

We have used an *in silico* mutational scan based on MutateX [41] and the FoldX energy function [42] to investigate whether AMBRA1 mutations validated for this study were likely to affect the stability of the β-propeller domain (**Fig. 6**). Our results show that out of the 7 experimentally validated mutations, 4 were mutational hotspots. These are residues 63, 110, 142 and 170 for which the substitution to most residue types was found destabilizing (ΔΔG >1.2 kcal/mol) (**Fig. 6**). On the contrary, the scan shows that any mutation at residue 90 was predicted not to affect stability, while residues 97 and 157 had a less extreme behavior, with only some substitutions having a negative effect (**Fig. 6**). Unsurprisingly, the experimentally tested mutations at the hotspot sites (P63S, L110F, S142F, P170S) were found to be destabilizing as well (**Fig. 6**). Mutations S90F and A157V had a neutral effect on stability, while T97I was classified as stabilizing (ΔΔG = −2.34 kcal/mol) (**Fig. 6**).

**Figure 6:**
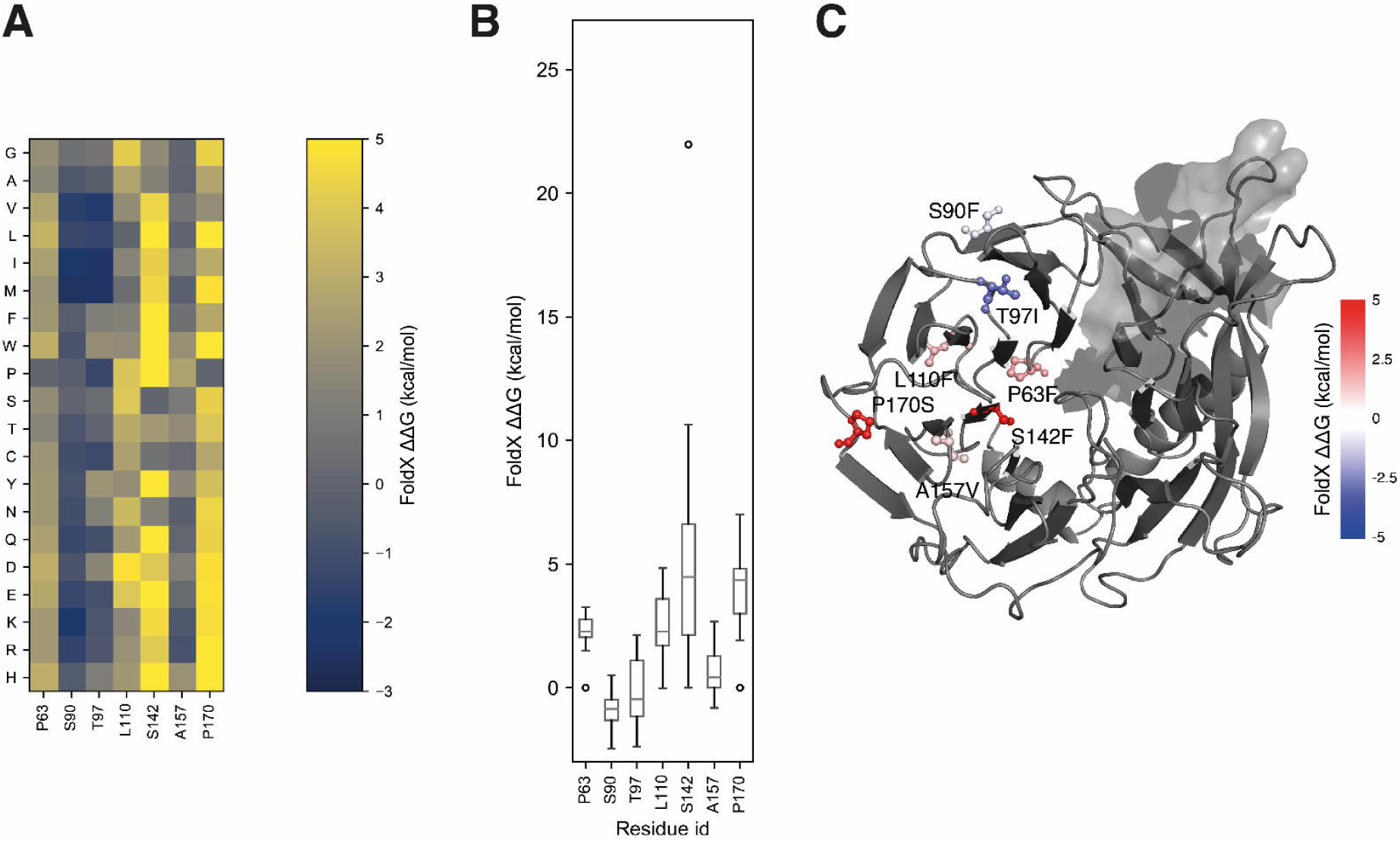
Predicted changes of folding free energy upon mutation for the mutations experimentally tested. In these plots, positive values represent mutations predicted to be destabilizing the protein structure, while negative values represent mutations predicted to be stabilizing. (**A**) Heatmap with predicted changes of free energy for the whole mutational scans at the sites we tested experimentally. Values in the plot have been limited within the −3 to 5 kcal/mol range to avoid using a less effective color scale due to outliers. (**B**) Box plots representing the distribution of the same values plotted in (**A**). (**C**) Residues at mutation sites corresponding to mutations that have been experimentally tested are shown on the predicted protein structure as sticks, colored proportionally to the predicted free energy change of the respective mutations, as per the colorbar. Values here have been limited in the −5 to 5 kcal/mol for the same reasons as in (**B**) to obtain a symmetrical colormap around 0.

## Discussion

In this work we have presented *Cancermuts*, a Python package for discovery, annotation and prioritization of cancer-related mutations. Our software can interrogate cancer genomics and mutation databases such as COSMIC and cBioPortal to retrieve cancer-associated mutations, both in pan-cancer or specific cancer types and studies. It also annotates both protein sequence and identified mutations to (i) give a context in terms of functional or structural features surrounding mutation sites and (ii) assess their potential to interfere with such features. Annotating mutations with pathogenicity scores such as REVEL and gnomAD allele frequencies also helps inform on their potential for pathogenicity more in general. *Cancermuts* has been designed as a Python package to ensure maximum flexibility, expandability and modularity. It is straightforward to use for people with basic Python skills and can be either used independently or incorporated in more complex workflows, e.g. after reducing the information it collects to a data frame.

In this contribution we have tested our approach on the protein AMBRA1, focusing on cancer mutations from melanoma. Indeed, melanoma is one of the cancer types in which AMBRA1 displays a crucial anti-tumorigenic role [24] and a mutational rate among the highest [21]. AMBRA1 is largely an intrinsically disordered protein (IDP), and its “unstructure” suggests that it can adapt to diverse situations and possibly coordinate different intracellular processes mainly by regulating protein-protein interactions [18]. The flexibility of its long disordered regions could play an important role in modulating the conformational changes needed to provide interaction surfaces that are complementary to different biological partners. Our tool identified several AMBRA1 mutations of potential interest in melanoma and allowed us to contextualize them in terms of localization in a predicted structured region, surrounding post-translational modifications, embedding in short linear motifs, and to annotate them for pathogenicity scores. Based on its importance in AMBRA1 itself [25] and Cyclin D1 [21-23] stability, we then focused our attention on the N-terminal region of the protein and assessed the effect of the most interesting mutations, given their context and annotations. Mutations have been selected by means of the pathogenicity score REVEL using a threshold of 0.4, which corresponds to a good balance between specificity and sensitivity (sensitivity 0.81 and specificity 0.85) [37] and represents a good compromise. Nonetheless, further fine-tuning of the cut-off might be necessary to suit different cases, also depending on the amount of available resources for further validation. Of the tested mutations, none had a clear effect on the AMBRA1-related autophagic or apoptotic pathways. However, we cannot rule out that such mutations might have effects we did not test for, or that such mutations might be detrimental in other conditions or cell types, or in conjunction with others. In this sense, having a wider range of readouts could help understand whether these false positive mutations can be important. This also highlights a potential downside of using pathogenicity predictors that are not tailored towards a specific disease or tissue. It has been shown that the performance of variant prioritization predictions varies with diseases phenotype [43], and machine learning models trained on more specific datasets could incorporate more of the cellular context of the identified variants or diseases, improving performance [43]. It should also be noted that all the N-terminal tested mutations fall in a generally very well conserved region of AMBRA1, as demonstrated by our protein sequence alignment among sequences of AMBRA1 orthologs from human, chimpanzee, mouse, rat, bovine, Xenopus and zebrafish (see **Methods** and **Table S4**). As 8 out of 18 of the pathogenicity scores integrated in REVEL are based on conservation, this is probably a contributing factor to the score that REVEL assigns to the mutations in this region. Interestingly, although differently, mutations of the conserved L110 (L→F) and P170 (P→S) residues were found implicated in functions of AMBRA1 recently reported to be relevant in terms of tumor growth and progression [21-24]. Among these, the expression of the L110F mutant (which maps next to the DDB1-Cullin4 interaction domain of AMBRA1), correlated with increased protein levels of Cyclin D1. Despite no difference in terms of proliferation was assessed (a counterintuitive outcome which may be explained by the high proliferative rate of A375 cells), the high Cyclin D1 levels detected in these circumstances may implicate an impaired ability of AMBRA1 to control Cyclin D1 stability. Moreover, both mutations increased the phosphorylation status of SRC (pSRC-Y416), a marker of the FAK1 signalling. Under the same conditions, cells expressing our mutants displayed higher cell invasiveness, hence suggesting a potential pathogenic effect for either mutation. Structure-based mutational scans suggest that both mutations are likely to destabilize the protein structure. Indeed, both positions were found to be extremely sensitive to mutation in a way that any amino acid change is likely to destabilize the protein at these positions. Despite the protein levels of the L110F and P170S mutants were not affected when screened using myc-tagged AMBRA1 mutants, this does not rule out that the protein structure itself might be affected –as suggested by the failure of detection by means of anti-AMBRA1 antibody, which was raised against residues 999-1298 of the AMBRA1 sequence, hence targeting the C-terminus. This includes part of the predicted β-propeller domain that bridges to the N-terminal region by means of a β-sheet. Residues 110 and 170 are not directly in contact with this region (**Fig. 6**), meaning it is unlikely their mutation would have a direct effect; nonetheless, they could still elicit a long-range effect by disrupting the local propeller structure and interfering with propeller assembly.

Even though these mutations feature the lowest REVEL score among those classified as pathogenic, they were found to bear the largest effects among those we tested. We speculated this might be due to their potential of inducing conformational changes or destabilization of the AMBRA1 protein structure. In this case, therefore, as REVEL does not include predictions of changes in protein structural stability explicitly, additional annotations that rely directly on structural information could complement and add another compelling layer of evidence. In this sense, tools able to perform high-throughput mutational scans (e.g. MutateX, which uses FoldX) aiming at predicting the impact of mutations on protein structure could be integrated in the *Cancermuts* package, for instance by including ready-made mutational scans in the annotation pipeline, which can be provided through the structure-based framework introduced in the work by Fas et al. [10].

*Cancermuts* was created with a modular design philosophy. This makes it possible to add additional layers of evidence by including support for them in the code, taking advantage of the pre-existing package structure. This will be useful to add new predictors or other data as they become available or to tailor its use to specific cases or datasets. For example, predicted structures from the AlphaFold protein structure models collection [44] could be used to integrate an additional layer of information about the structure and differentiate between predicted disordered and ordered regions. Other attractive layers of evidence also include predictors for the effect of mutation based on sequence evolution, such as the recently released EVE model [45] which relies on generative models of evolutionary data, GEMME [46], DeepSequence [47] or EVmutation [48]. The fact that *Cancermuts* also allows user-curated input yet adds another layer of flexibility, allowing to add information without need to write any code.

## Materials and Methods

### Cell Lines and Treatments

The human melanoma cell line A375 (RRID: CVCL_0132) was cultivated in GlutaMAX™-additioned Dulbecco’s Modified Eagle Medium (DMEM) (ThermoFisher Scientific; cat# 31966-021) supplemented with 10% FBS (ThermoFisher Scientific; cat# 10270-106) and 100 U/ml P/S (ThermoFisher Scientific; cat# 15140122). Cells were cultured in an atmosphere of 5% CO_2_ in air at 37°C and passaged no more than 15 times. Cells were used within a few months of resuscitation and routinely tested for *Mycoplasma* during sub-cultivation by PCR-based methods (eurofins Genomics, DE) and only used if negative. During the experiments, cells were plated at a density of 1×10^5^ cells/ml, unless otherwise indicated. Chloroquine (CQ, Sigma-Aldrich; cat# C6628) and MG-132 (Sigma-Aldrich; cat# M7449) were dissolved in DMSO and used at final concentrations of 40 µM and 10µM, respectively, for 4h while DMSO was used in control cells.

### siRNAs and Transfection Methods

Reverse siRNA transfection was performed at the time of seeding at a final 20 nM concentration for a total of 48h, unless otherwise indicated. siRNA sequences for AMBRA1 are custom designed, as reported in [24]. Negative control cells (siScr) were transfected with the MISSION^®^ siRNA Universal Negative Control #1 (Sigma-Aldrich; cat# SIC001). Overexpression of plasmid constructs was carried for the times indicated in the specific experiments after cells had been reversely transfected for 24h with siAMBRA#2, which was specifically designed to target the 5’-UTR region of *AMBRA1* in order to exclude effect(s) (i) of the siRNA on expression of AMBRA1 plasmid constructs and (ii) of endogenous AMBRA1 in the downstream applications (**Fig. 3B**). The sequence coding for wild-type AMBRA1 (WT) (UniProt ID: Q9C0C7-1) was cloned in either pcDNA™3.1 Mammalian Expression (ThermoFisher Scientific; cat# V79020) or pcDNA™3.1/myc-His A, B, & C Mammalian Expression (ThermoFisher Scientific; cat# V80020) vectors. The coding sequences were amplified by PCR and cloned in the acceptor vector by means of EcoRI and NotI restriction sites. AMBRA1 mutants (P63S, S90F, T97I, L110F, S142F, A157V, P170S) were generated by site-directed mutagenesis using AMBRA1 as template and custom designed primers. The list of mutants of the N-terminal region of AMBRA1 on which the *in vitro* validation has been performed does not include all the point mutations with REVEL ≥ 0.4 shown in **Table S2**, as DNA constructs were generated on a previous version of the mutation plot dated May 22^nd^, 2018. All transfections were performed using Lipofectamine™ 2000 Transfection Reagent (ThermoFisher Scientific; cat# 11668-019), and manufacturer’s instructions were followed.

### Protein Expression Analysis

At the time of collection, cells were washed in Phosphate Buffer Solution (PBS, ThermoFisher Scientific; cat# 14190144), mechanically detached and centrifuged at 1,200×g for 5 min at 4°C and cell pellets processed and previously described [24]. Protein concentration of supernatants was determined by the Lowry’s method. For soluble/insoluble analysis, cell debris (insoluble fraction) was washed three times in RIPA buffer followed by centrifugation at 10,000×g for 5 min at 4°C. Both soluble and insoluble fractions were denatured in NuPAGE™ LDS Sample Buffer (4X) (ThermoFisher Scientific; cat# NP0007) supplemented with NuPAGE™ Sample Reducing Agent (10X) (ThermoFisher Scientific; cat# NP0004) followed by incubation at 100°C for 5 min. Protein extracts were separated by SDS-PAGE using Criterion™ TGX™ Precast Gels (Bio-Rad Laboratories; cat# 567-8084) and blotted onto PVDF membranes (Bio-Rad Laboratories; cat# 10026933) using a Trans-Blot® Turbo™ Transfer System (Bio-Rad Laboratories). Primary antibodies used are as follows:

| Target | Dilution | Distributor | Cat# | RRID: |
| --- | --- | --- | --- | --- |
| Actin | 1:40,000 | Novus Biologicals | NB600-501 | AB_10077656 |
| AMBRA1 | 1:1,000 | Santa Cruz Biotechnology | sc-398204 | / |
| c-myc | 1:1,000 | Santa Cruz Biotechnology | sc-40 | / |
| CASP-3 | 1:1,000 | Cell Signaling Technology | 9662S | AB_331439 |
| Cyclin D1 | 1:2,000 | Abcam | ab16663 | AB_443423 |
| LC3 | 1:2,500 | Cell Signaling Technology | 3868S | AB_2137707 |
| pSRC-Y416 | 1:1,000 | Cell Signaling Technology | 6943S | AB_10013641 |
| SRC | 1:1,000 | Cell Signaling Technology | 2123S | AB_2106047 |
| Ubiquitin (Ub) | 1:1,000 | Merck-Millipore | 05-1308 | AB_1587580 |

Images were captured with a ChemiDoc™ MP System (Bio-Rad Laboratories; cat# 1708-280) provided with the Image Lab 6.0.1 Software (Bio-Rad Laboratories). Densitometry analyses were carried out using the ImageJ software (1.52.q) (University of Wisconsin; RRID:SCR_003070).

### RNA Isolation, Reverse Transcription and Quantitative RT-PCR

Total RNA was isolated and reverse transcription were performed as previously described [24]. cDNA was diluted 3 times and mRNA expression levels detected by PowerUp™ SYBR™ Green Master Mix (ThermoFisher Scientific; cat# A25742), according to the instructions, on a ViiA 7 Real-Time PCR System v1.3 (Applied Biosystems). All reactions were run in triplicate and mRNA levels expressed as fold change (relative to control) after normalization to the internal housekeeping *L34*. The specific primer pairs were custom designed and tested with Primer-BLAST (NCBI; RRID:SCR_003095). Primers used were obtained from TAG Copenhagen A/S (Copenhagen, DK) and are as follows: *L34*: FW: 5’-GGC CCT GCT GAC ATG TTT CTT −3’, RV: 5’-GTC CCG AAC CCC TGG TAA TAG A −3’; *AMBRA1*: FW: 5’-AAC CCT CCA CTG CGA GTT GA −3’, RV: 5’-TCT ACC TGT TCC GTG GTT CTC −3’; *FN1*: FW: 5’-CGA CAC ATT CCA CAA GCG TC −3’, RV: 5’-CAT TGG TCG ACG GGA TCA CA −3’; *VIM*: FW: 5’-GAC GCC ATC AAC ACC GAG TT −3’, RV: 5’-CTT TGT CGT TGG TTA GCT GGT −3’.

### Wound Healing Assay

24h after re-expression of the plasmid constructs, 25,000 cells were seeded in each of the 2 wells of silicone inserts with a defined gap of 500 µm (ibidi®; cat# 80209) in 6-well plates. After 16h, the inserts were removed and wound closure followed at the times indicated. Migrating cells were imaged with an IX71 inverted microscope (Olympus) provided with a CellSens Imaging Software 2 (Olympus; RRID:SCR_016238). The area of wound closure was calculated using ImageJ with respect to the initial area (T_0_) and expressed as percentage of wound healing at the time points indicated. In the representative pictures, the white and yellow lines outline the edge of the wound at T_0_ and at 24h, respectively.

### Cell proliferation

24h after re-expression of the plasmid constructs, 10,000 cells were seeded in 12-well plates. After 24h and 48h, cells were washed with PBS, fixed-and-stained with a 0.025% (w/v) Crystal violet (Sigma-Aldrich; cat# C6158) solution in 20% (v/v) MeOH on ice for 15 minutes. After washing with ddH_2_O, plates were air-dried and pictures taken with an IX71 inverted microscope (Olympus) provided with a CellSens Imaging Software 2 (Olympus). For quantitation, Crystal violet was eluted with 100% MeOH and absorbance measured at 595 nm by a VICTOR Multilabel Plate Reader (PerkinElmer). Data are expressed as fold change with respect to absorbance of control sample (WT at 24h).

### Cell viability

24h after re-expression of the plasmid constructs, 7,500 cells were seeded in 96-well plates and cell viability measured at the times indicated using the Cell Counting Kit-8 (Dojindo; cat# CK04-11) at 450□nm using a VICTOR Multilabel Plate Reader (PerkinElmer) after 2□h of incubation, according to the manufacturer’s instructions. Data are shown as fold change of viable cells with respect to control cells (WT at 24h).

### Statistical analysis

Ordinary one-way ANOVA was used for densitometry and RT-qPCR analyses. Two-way ANOVA was used for wound healing, cell proliferation and viability assays. All ANOVA tests were corrected using the Bonferroni multiple comparison test and statistical values calculated in function of a control case. GraphPad/Prism9 (version 9.2.0) (RRID:SCR_002798) was used for plotting graphs and to perform statistical analysis. Data are presented as means ± SEM or SD, as indicated in the figure legends, and significance was designated as follows: *p□≤ □0.05; **p□≤ □0.01; ***p□≤ □0.001; ****p□≤ □0.0001; ns, not significant.

### Structured regions of AMBRA1 according to AlphaFold

We have downloaded the Alphafold [49] model for human AMBRA1 from the EMBL-EBI Alphafold Protein Structure Database [44], entry Q9C0C7. Visual inspection of the model showed a major structural feature for this model – a β-propeller folded domain spanning regions ∼41-203 and ∼857-1040 of AMBRA1. The AlphaFold prediction was deemed to be confident (pLDDT > 70) for the first stretch of residues and for most part of the second, with short stretches of residues at lower confidence which correspond to short solvent-exposed loops and are thus likely to be disordered. AlphaFold also predicts the N-terminus of AMBRA1 to be structured as two consecutive alpha helices, one with low confidence (residues 7-19, most of them with pLDDT scores between 50 and 70) and one at high confidence (residues 25-40, pLDDT > 70)

### Free energy calculations

We trimmed the structure keeping only residue stretches corresponding to the predicted structured regions of AMBRA1 (residues 1-200 and 850-1040). We then used the MutateX pipeline [41] saturation scan protocol with FoldX 5.0 [42] to run a complete mutational scan of the resulting structure, predicting the changes of folding free energy upon mutation for the substitution of each amino acid to each natural variant for a total of 7820 data points. For each data point we considered the average difference in free energy between wild-type and mutant variant over five independent FoldX runs. The MutateX protocol includes both a repair step, in which the structure is optimized using FoldX, and generation of mutant variant structures together with folding free energy estimation.

### Sequence alignment

We have obtained a protein multiple sequence alignment between different AMBRA1 orthologs using Clustal Omega [50], using the protein sequences corresponding to the main protein isoform of AMBRA1 of human, chimpanzee, mouse, rat, bovine, *Xenopus* and zebrafish (ambra1a for the latter).

## Acknowledgments

The authors would like to particularly thank Vanda Turcanova for her technical support and help with cloning, and Laila Fischer for help with secretarial work.

## Author contributions

Conception and design: M.T., L.D.L., D.D.Z., E.P.; Development of methodology: M.T., L.D.L., D.D.Z., E.P.; Acquisition of data: M.T., L.D.L., M.V.V., R.K., E.P.; Analysis and interpretation of data: M.T., L.D.L., D.D.Z., E.P.; Writing, review and/or revision of the manuscript: M.T., L.D.L., F.C., D.D.Z., E.P. (all authors have reviewed and edited the manuscript); Administrative, technical, or material support: M.T., L.D.L., F.C., D.D.Z., E.P.; Study supervision: D.D.Z. and E.P..

## Ethics statement

No human participants, human tissue, or animal model are involved in this study.

## Additional Information

Supplementary Fig. 1.

Table S1

Table S2

Table S3

Table S4

