## Supplementary Figure 1 for "The *Cancermuts* software package for the prioritization of missense cancer variants: a case study of AMBRA1 in melanoma"

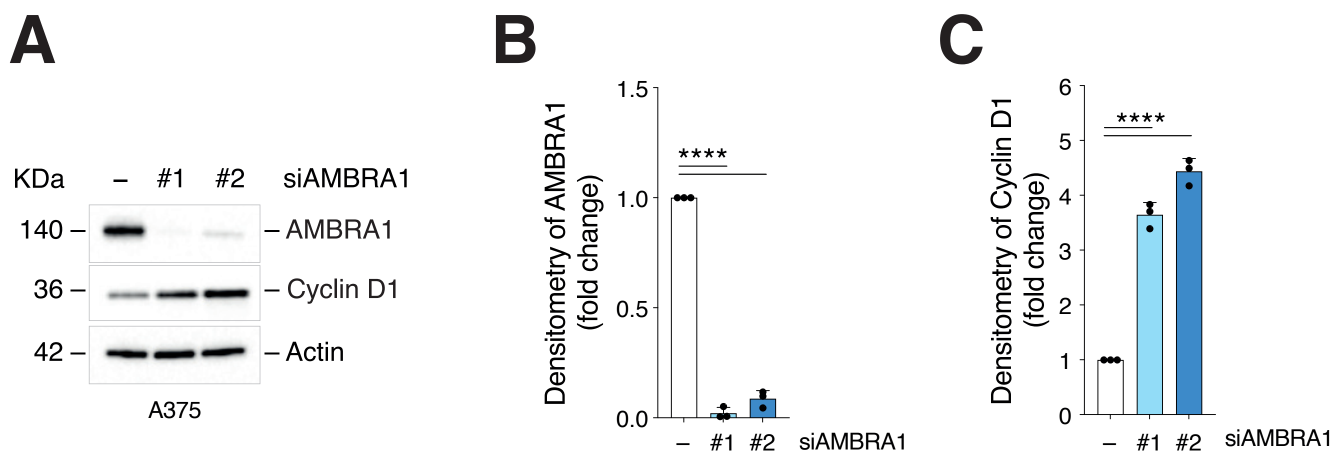


**Supplementary Figure 1. Expression of Cyclin D1 upon AMBRA1 silencing in melanoma cells.** (**A**) A375 cells were silenced for *AMBRA1* (siAMBRA1#1 and #2) and WB analyses performed to detect Cyclin D1 levels. AMBRA1 and Actin were used as transfection and loading control, respectively. Images are representative of n=3 independent experiments and are quantified in (**B**) for AMBRA1 and in (**C**) for Cyclin D1. Data are expressed as fold change ± SD vs control cells after normalization on Actin (n=3; ****$p<0.0001$; one-way ANOVA).
