## Supplemental Table S4 for "The *Cancermuts* software package for the prioritization of missense cancer variants: a case study of AMBRA1 in melanoma"

multiple sequence alignment between the protein sequence of different AMBRA1 orthologs using Clustal Omega [Madeira, F., et al., *Search and sequence analysis tools services from EMBL-EBI in 2022.* Nucleic Acids Res, 2022.], using the protein sequences corresponding to the main Uniprot protein isoform of AMBRA1 from human (AMRA1_HUMAN), chimpanzee (A0A2I3TB21_PANTR), mouse (AMRA1_MOUSE), rat (F1LNQ2_RAT), bovine (F1MPW0_BOVIN), *Xenopus* (F6YH53_XENTR) and zebrafish ambra1a (AMR1A_DANRE).

```
CLUSTAL O(1.2.4) multiple sequence alignment


sp|E7FAG6|AMR1A_DANRE                   -MKLGQRNSVCILSSRERGAPGLASYRVLQQLVEEKTQRMKWQSQKVELPDSPRSTFLLA  59
tr|F6YH53|F6YH53_XENTR                  MKVVPEKNAVRILAGRERGLQAQGAQRLLQCLVEDKTRFMKWEGKKVELPDSPRSTFLLA  60
tr|F1MPW0|F1MPW0_BOVIN                  MKVVPEKNAVRILWGRERGTRTFGAQRLLQELVEDKTRCMKWEGKRVELPDSPRSTFLLA  60
sp|Q9C0C7|AMRA1_HUMAN                   MKVVPEKNAVRILWGRERGARAMGAQRLLQELVEDKTRWMKWEGKRVELPDSPRSTFLLA  60
tr|A0A2I3TB21|A0A2I3TB21_PANTR          MKVVPEKNAVRILWGRERGARAMGAQRLLQELVEDKTRWMKWEGKRVELPDSPRSTFLLA  60
sp|A2AH22|AMRA1_MOUSE                   MKVVPEKNAVRILWGRERGTRAMGAQRLLQELVEDKTRWMKWEGKRVELPDSPRSTFLLA  60
tr|F1LNQ2|F1LNQ2_RAT                    MKVVPEKNAVRILWGRERGTRAMGAQRLLQELVEDKTRWMKWEGKRVELPDSPTLDFLLA  60
                                           : ::*:* ** .****    .: *:** ***:**: ***:.::*******   ****

sp|E7FAG6|AMR1A_DANRE                   FSPDRSLMASTHVNHNIYITEVKSGKCVHSLVGHRRTPWCLTFHPIIPGLIASGCLDGEV  119
tr|F6YH53|F6YH53_XENTR                  FSPDRTLLASTHVNHNIYITEVKTGRCIHSLVGHRRTPWCVTFHPTIPGLIASGCLDGEV  120
tr|F1MPW0|F1MPW0_BOVIN                  FSPDRTLLASTHVNHNIYITEVKTGKCVHSLIGHRRTPWCVTFHPTISGLIASGCLDGEV  120
sp|Q9C0C7|AMRA1_HUMAN                   FSPDRTLLASTHVNHNIYITEVKTGKCVHSLIGHRRTPWCVTFHPTISGLIASGCLDGEV  120
tr|A0A2I3TB21|A0A2I3TB21_PANTR          FSPDRTLLASTHVNHNIYITEVKTGKCVHSLIGHRRTPWCVTFHPTISGLIASGCLDGEV  120
sp|A2AH22|AMRA1_MOUSE                   FSPDRTLLASTHVNHNIYITEVKTGKCVHSLIGHRRTPWCVTFHPTISGLIASGCLDGEV  120
tr|F1LNQ2|F1LNQ2_RAT                    FSPDRTLLASTHVNHNIYITEVKTGKCVHSLIGHRRTPWCVTFHPTISGLIASGCLDGEV  120
                                        *****:*:***************:*:*:***:********:**** * ************

sp|E7FAG6|AMR1A_DANRE                   RIWDLHGGSESWLTESNSAIASLAFHPTAQLLLIATNNEVHLWDWSRKEPFTVVKTASET  179
tr|F6YH53|F6YH53_XENTR                  RIWDLHGGSESWFTDSNNAIASLAFHPTVQLLLIATASEIHFWDWSRREPFAVVKTASEM  180
tr|F1MPW0|F1MPW0_BOVIN                  RIWDLHGGSESWFTDSNNAIASLAFHPTAQLLLIATASEIHFWDWSRREPFAVVKTASEM  180
sp|Q9C0C7|AMRA1_HUMAN                   RIWDLHGGSESWFTDSNNAIASLAFHPTAQLLLIATANEIHFWDWSRREPFAVVKTASEM  180
tr|A0A2I3TB21|A0A2I3TB21_PANTR          RIWDLHGGSESWFTDSNNAIASLAFHPTAQLLLIATANEIHFWDWSRREPFAVVKTASEM  180
sp|A2AH22|AMRA1_MOUSE                   RIWDLHGGSESWFTDSNNAIASLAFHPTAQLLLIATANEIHFWDWSRREPFAVVKTASEM  180
tr|F1LNQ2|F1LNQ2_RAT                    RIWDLHGGSESWFTDSNNAIASLAFHPTAQLLLIATANEIHFWDWSRREPFAVVKTASEM  180
                                        ************:*:**.**********.******* .*:*:*****:***:*******

sp|E7FAG6|AMR1A_DANRE                   ERVRLVRFDPLGHYLLTAIVNPSNQPNDDDPEIPMDSVEMPHLRQRSFLQSQPARRTPIL  239
tr|F6YH53|F6YH53_XENTR                  ERVRLVRFDPLGHYLLTAIVNPSNQQSDEESEIPLDSAEIPIYRQRSLLQTQPVRRTPLL  240
tr|F1MPW0|F1MPW0_BOVIN                  ERVRLVRFDPLGHYLLTAIVNPSNQQGDDEPEIPVDGAELSHYRQRALLQSQPVRRTPLL  240
sp|Q9C0C7|AMRA1_HUMAN                   ERVRLVRFDPLGHYLLTAIVNPSNQQGDDEPEIPIDGTELSHYRQRALLQSQPVRRTPLL  240
tr|A0A2I3TB21|A0A2I3TB21_PANTR          ERVRLVRFDPLGHYLLTAIVNPSNQQGDDEPEIPIDGTELSHYRQRALLQSQPVRRTPLL  240
sp|A2AH22|AMRA1_MOUSE                   ERVRLVRFDPLGHYLLTAIVNPSNQQGDDEPEIPIDGTELSHYRQRALLQSQPVRRTPLL  240
tr|F1LNQ2|F1LNQ2_RAT                    ERVRLVRFDPLGHYLLTAIVNPSNQQGDDEPEIPIDGTELSHYRQRALLQSQPVRRTPLL  240
                                        ************************* .*:: ***:*..*:   ***::**:**.****:*

sp|E7FAG6|AMR1A_DANRE                   HNFLHILTSRNSVPQAGGAHSASTDG---------------------------SSDSSG  271
tr|F6YH53|F6YH53_XENTR                  HNFLHMLSSRSSGTQAGEQPPVQDSATPSPPPPPPPPLPPPSNDSNQRVLYTRIRERS-P  299
tr|F1MPW0|F1MPW0_BOVIN                  HNFLHMLSSRSSCIQVGEQSTVQDPATPSPPPPPPPQPST----ERPRTSAYIRLRQRVSY  296
sp|Q9C0C7|AMRA1_HUMAN                   HNFLHMLSSRSSGIQVGEQSTVQDSATPSPPPPPPPQPST----ERPRTSAYIRLRQRVSY  296
tr|A0A2I3TB21|A0A2I3TB21_PANTR          HNFLHMLSSRSSGIQVGEQSTVQDSATPSPPPPPPPQPST----ERPRTSTYIRLRQRVSY  296
sp|A2AH22|AMRA1_MOUSE                   HNFLHMLSSRSSGIQVGEQSTVQDSATPSPPPPPPPQPST----ERPRTSAYIRLRQRVSY  296
tr|F1LNQ2|F1LNQ2_RAT                    HNFLHMLSSRSSGIQVGEQSTVQDSATPSPPPPPPPQPST----ERPRTSAYIRLRQRVSY  296
                                        *****:*:**.*  *.*    ..  .                              .

sp|E7FAG6|AMR1A_DANRE                   PYTLMCVQPLGMVCFCSRCSAARVPSPPDEDPS----DSASLEAQAHTFSSARTEPLQMS  327
tr|F6YH53|F6YH53_XENTR                  GVSTPCCQDPSMLCLCIRCSSARLSSPLFPHPEAPGAPSTSSGVTSTSFSPVQTEPSHSL  359
tr|F1MPW0|F1MPW0_BOVIN                  PT-AECCQHLGILCLCSRCSGTRVPSLL-PHQD--SVPPASARATTPSFSFVQTEPFHPP  352
sp|Q9C0C7|AMRA1_HUMAN                   PT-AECCQHLGILCLCSRCSGTRVPSLL-PHQD--SVPPASARATTPSFSFVQTEPFHPP  352
tr|A0A2I3TB21|A0A2I3TB21_PANTR          PT-AECCQHLGILCLCSRCSGTRVPSLL-PHQD--SVPPASARATTPSFSFVQTEPFHPP  352
sp|A2AH22|AMRA1_MOUSE                   PTTVECCQHPGILCLCSRCAGTRVPSLL-PHQD--SVPPASARATTPSFSFVQTEPFHPP  353
tr|F1LNQ2|F1LNQ2_RAT                    PTTVECCQHPGILCLCSRCAGTRVPSLL-PHQD--SVPPASARATTPSFSFVQTEPFHPP  353
                                             * *  .::*:* **:.:*: *     . .       :*  . : :** .:*** :

sp|E7FAG6|AMR1A_DANRE                   R---------FSVESRAANRSSAFSSVYG--GGSNMRNHSSSSGRRGVTGMAPVPHFRQH  376
tr|F6YH53|F6YH53_XENTR                  ERQQQQHQSSAQHDHGLLSRPSAFSSVHSSTAGNTLRNLSLGPTRRSLTG----QPSRY-  414
tr|F1MPW0|F1MPW0_BOVIN                  E-----QASSAPHDPGLLSRPSAFSTVQSSTAGNTLRNLSLGPPRRSLGGPLSGHPSRY-  406
sp|Q9C0C7|AMRA1_HUMAN                   E-----QASSTQQDQGLLNRPSAFSTVQSSTAGNTLRNLSLGPTRRSLGGPLSSHPSRY-  406
tr|A0A2I3TB21|A0A2I3TB21_PANTR          E-----QASSTQQDQGLLNRPSAFSTVQSSTAGNTLRNLSLGPTRRSLGGPLSSHPSRY-  406
sp|A2AH22|AMRA1_MOUSE                   E-----QASSTQQDQGLLNRPSAFSTVQSSTAGNTLRNLSLGPTRRSLGGPLSSHPSRY-  407
tr|F1LNQ2|F1LNQ2_RAT                    E-----QASSTQQDQGLLNRPSAFSTVQSSTAGNTLRNLSLGPTRRSLGGPLSSHPSRY-  407
                                        .             :    .* ****:* .  .*..:** * .  **.: *       *

sp|E7FAG6|AMR1A_DANRE                   PPGREGGGRHPGADWTVSGL----NGQSSSMTPQRTGASSVSLLSVLRQQETSFQSPVYT  432
tr|F6YH53|F6YH53_XENTR                  --QQ-PSTDLAGSEWTRTVLSMGPRPEMEPMPPPRTSASSVSILSVLRQQEGETSSSVYT  471
tr|F1MPW0|F1MPW0_BOVIN                  --HRDIASGLTGSEWTRTVLSLNSRSEAESMPPPRTSASSVSLLSVLRQQEGGSQASVYT  464
sp|Q9C0C7|AMRA1_HUMAN                   --HREIAPGLTGSEWTRTVLSLNSRSEAESMPPPRTSASSVSLLSVLRQQEGGSQASVYT  464
tr|A0A2I3TB21|A0A2I3TB21_PANTR          --HREIAPGLTGSEWTRTVLSLNSRSEAESMPPPRTSASSVSLLSVLRQQEGGSQASVYT  464
sp|A2AH22|AMRA1_MOUSE                   --HRELAPGLTGSEWTRTVLTLNSRSEVESMPPPRTSASSVSLLSVLRQQEGGSQASVYT  465
tr|F1LNQ2|F1LNQ2_RAT                    --HRELAPGLTGSEWTRTVLTLNSRSEVESMPPPRTSASSVSLLSVLRQQEGGSQASVYT  465
                                          :  .    *::** : *    . : . * * **.*****:********   .: ***
```

```
sp|E7FAG6|AMR1A_DANRE               SASDRWGSTPGTSSSRHRPPEEEGQSSSSSIHSVLRCNLYRYFMDYEGTQDTVQPLDGSR  492
tr|F6YH53|F6YH53_XENTR              SATEGRGFSSSES----DSSAAPPPVHPTTTRTELQCDLRRFFLEYDRLHELEPGAPGTT  527
tr|F1MPW0|F1MPW0_BOVIN              SATEGRGFPASGLAAESDGGNGSSQNNSGSIRHELQCDLRRFFLEYDRLQELDQSLSGEA  524
sp|Q9C0C7|AMRA1_HUMAN               SATEGRGFPASGLATESDGGNGSSQNNSGSIRHELQCDLRRFFLEYDRLQELDQSLSGEA  524
tr|A0A2I3TB21|A0A2I3TB21_PANTR      SATEGRGFPASGLATESDGGNGSSQNNSGSIRHELQCDLRRFFLEYDRLQELDQSLSGEA  524
sp|A2AH22|AMRA1_MOUSE               SATEGRGFPSSGLATESDGGNGSSQNNSGSIRHELQCDLRRFFLEYDRLQELDQSLSGET  525
tr|F1LNQ2|F1LNQ2_RAT                SATEGRGFPSSGLATESDGGNGSSQNNSGNIRHELQCDLRRFFLEYDRLQELDQSLSGET  525
                                    **::  *   .                  . :  *:*:* *:*::*:  ::      *

sp|E7FAG6|AMR1A_DANRE               -QDQQTQEMLNNNMDPEQPGPSHYQS----PYSGENPPHSHMNRCRVCHNLFTYNQGSRR  547
tr|F6YH53|F6YH53_XENTR              --QGQTHEMLNNNLEPEQPGPSHQPPQAHSGDSTSNQPRGHINRCRACHNLLTFNNDTLR  585
tr|F1MPW0|F1MPW0_BOVIN              PQAQQAQEMLNNNLESERPGPSHQPT-PHSSENNSNLSRGHLNRCRACHNLLTFNNDTLR  583
sp|Q9C0C7|AMRA1_HUMAN               PQTQQAQEMLNNNIESERPGPSHQPT-PHSSENNSNLSRGHLNRCRACHNLLTFNNDTLR  583
tr|A0A2I3TB21|A0A2I3TB21_PANTR      PQTQQAQEMLNNNIESERPGPSHQPT-PHSSENNSNLSRGHLNRCRACHNLLTFNNDTLR  583
sp|A2AH22|AMRA1_MOUSE               PQTQQAQEMLNNNIESERPGPSHLPT-PHSSENNSNLSRGHLNRCRACHNLLTFNNDTLR  584
tr|F1LNQ2|F1LNQ2_RAT                PQTQQAQEMLNNNIESERPGPSHQPT-PHSSENNSNLSRGHLNRCRACHNLLTFNNDTLR  584
                                        *::******:: *:*****         . .*  :.*:****.****:*:*:.: *

sp|E7FAG6|AMR1A_DANRE               WDRTGQPS-STERNTPWQPSSSAFHSVA--PVSQSNEHLLEHRPIESTPNT--PEPHVPF  602
tr|F6YH53|F6YH53_XENTR              WERTPPTYPETAGTSTWQPPPIPPPSSFQAGLAAGSTDHPGTPRVERVDLGSMAASSNRL  645
tr|F1MPW0|F1MPW0_BOVIN              WERSTPNYSSGEASASWQVPGTF-----E-GMAASGSQLP---PLERTE-GQ-TASSSRL  632
sp|Q9C0C7|AMRA1_HUMAN               WERTTPNYSSGEASSSWQVPSSF-----E-SVPSSGSQLP---PLERTE-GQ-TPSSSRL  632
tr|A0A2I3TB21|A0A2I3TB21_PANTR      WERTTPNYSSGEASSSWQVPSSF-----E-SVPSSGSQLP---PLERTE-GQ-TPSSSRL  632
sp|A2AH22|AMRA1_MOUSE               WERTTPNYSSGEASSSWHVSTTF-----E-GMPPSGNQLP---PLERTE-GQ-MPSSSRL  633
tr|F1LNQ2|F1LNQ2_RAT                WERTTPNYSSGEASSSWHVSTTF-----E-GMPPSGNQLP---PLERTE-GQ-MPNSSRL  633
                                    *:*:     .   .: *:             :  .. .      :* .           :

sp|E7FAG6|AMR1A_DANRE               SQRTDTGQHEEQAVGLVFNQETGQLERVYRQSAS-SRSANISQGALNQEMPEDTPDNDYL  661
tr|F6YH53|F6YH53_XENTR              ---ENGTTQEERTVGVVYNPETGHWERVYSQPASVSRAPNVSQEALPQDLQEESTEEDSL  702
tr|F1MPW0|F1MPW0_BOVIN              ELGNSASPQEERTVGVAFNQETGHWERIYTQA---SRSGTVSQEALHQDLPEESSEEDSL  689
sp|Q9C0C7|AMRA1_HUMAN               ELSSSASPQEERTVGVAFNQETGHWERIYTQS---SRSGTVSQEALHQDMPEESSEEDSL  689
tr|A0A2I3TB21|A0A2I3TB21_PANTR      ELSSSASPQEERTVGVAFNQETGHWERIYTQS---SRSGTVSQEALHQDMPEESSEEDSL  689
sp|A2AH22|AMRA1_MOUSE               ELSSSASSQEERTVGVAFNQETGHWERIYTQS---SRSGTVSQEALHQDMPEESSEEDSL  690
tr|F1LNQ2|F1LNQ2_RAT                ELSSSASSQEERTVGVAFNQETGHWERIYTQS---SRSGTVSQEALHQDMPEESSEEDSL  690
                                        .   :**::**:.:* ***: **:* *    **: .:** ** *:: *:: ::* *

sp|E7FAG6|AMR1A_DANRE               R----------------------------RLSPAAYYAQRMIQYLSRRDSVRQHSHRPP  692
tr|F6YH53|F6YH53_XENTR              RRRLLESSLISLSRYDATGSREHPIYPDPARLSPAAYYAQRMIQYLSRRDSIRQRSMRYQ  762
tr|F1MPW0|F1MPW0_BOVIN              R----------------------------RLSPAAYYAQRMIQYLSRRDSIRQRSMRYQ  720
sp|Q9C0C7|AMRA1_HUMAN               RRRLLESSLISLSRYDGAGSREHPIYPDPARLSPAAYYAQRMIQYLSRRDSIRQRSMRYQ  749
tr|A0A2I3TB21|A0A2I3TB21_PANTR      RRRLLESSLISLSRYDGAGSREHPIYPDPARLSPAAYYAQRMIQYLSRRDSIRQRSMRYQ  749
sp|A2AH22|AMRA1_MOUSE               RRRLLESSLISLSRYDGAGSREHPIYPDPARLSPAAYYAQRMIQYLSRRDSIRQRSMRYQ  750
tr|F1LNQ2|F1LNQ2_RAT                RRRLLESSLISLSRYDGAGSREHPIYPDPARLSPAAYYAQRMIQYLSRRDSIRQRSMRYQ  750
                                    *                             ********************:**:* *

sp|E7FAG6|AMR1A_DANRE               SRPRPLSSNPSSLSPSPVPNAESSEVDFEEFEENGSR--YRTPRNARMSAPSLGRFVGTR  750
tr|F6YH53|F6YH53_XENTR              QNR-LRTSAVSSSSEAQGA-VETGDLEYEEFEDNGDRSRHRTTRNARMSAPSLGRFVP-R  819
tr|F1MPW0|F1MPW0_BOVIN              QNR-LRSSTSSSSSDNQGPTVEGTDLEFEDFEDSGDRSRHRAPRNARMSAPSLGRFVP-R  778
sp|Q9C0C7|AMRA1_HUMAN               QNR-LRSSTSSSSSDNQGPSVEGTDLEFEDFEDNGDRSRHRAPRNARMSAPSLGRFVP-R  807
tr|A0A2I3TB21|A0A2I3TB21_PANTR      QNR-LRSSTSSSSSDNQGPSVEGTDLEFEDFEDNGDRSRHRAPRNARMSAPSLGRFVP-R  807
sp|A2AH22|AMRA1_MOUSE               QNR-LRSSTSSSSSDNQGPSVEGTDLEFEDFEDNGDRSRHRAPRNARMSAPSLGRFVP-R  808
tr|F1LNQ2|F1LNQ2_RAT                QNR-LRSSTSSSSSDNQGPSVEGTDLEFEDFEDNGDRSRHRAPRNARMSAPSLGRFVP-R  808
                                    ..     :*  ** *      .*  ::::*:**:.*.*  :*: *************  *

sp|E7FAG6|AMR1A_DANRE               RFLLPEFLPYAGIFHERGQPGLATHSSVNRVLAGAVIGDGQSAVASNIANTTYRLQWWDF  810
tr|F6YH53|F6YH53_XENTR              RFLLPEYLPYAGIFHERGQPGLATHSSVNRVLAGAVIGDGQSAVASNIANTTYRLQWWDF  879
tr|F1MPW0|F1MPW0_BOVIN              RFLLPEYLPYAGIFHERGQPGLATHSSVNRVLAGAVIGDGQSAVASNIANTTYRLQWWDF  838
sp|Q9C0C7|AMRA1_HUMAN               RFLLPEYLPYAGIFHERGQPGLATHSSVNRVLAGAVIGDGQSAVASNIANTTYRLQWWDF  867
tr|A0A2I3TB21|A0A2I3TB21_PANTR      RFLLPEYLPYAGIFHERGQPGLATHSSVNRVLAGAVIGDGQSAVASNIANTTYRLQWWDF  867
sp|A2AH22|AMRA1_MOUSE               RFLLPEYLPYAGIFHERGQPGLATHSSVNRVLAGAVIGDGQSAVASNIANTTYRLQWWDF  868
tr|F1LNQ2|F1LNQ2_RAT                RFLLPEYLPYAGIFHERGQPGLATHSSVNRVLAGAVIGDGQSAVASNIANTTYRLQWWDF  868
                                    ******:*****************************************************

sp|E7FAG6|AMR1A_DANRE               TKFDLPEISNASVNVLVPNCKIYNDASCDISADGQLLAVFIPSSQRGFPDEGILAVYSLA  870
tr|F6YH53|F6YH53_XENTR              TKYDLPEISNASINVLVQNCKIYNDASCDISADGQLLAAFIPSSQRGFPDEGILAVYSLA  939
tr|F1MPW0|F1MPW0_BOVIN              TKFDLPEISNASVNVLVQNCKIYNDASCDISADGQLLAAFIPSSQRGFPDEGILAVYSLA  898
sp|Q9C0C7|AMRA1_HUMAN               TKFDLPEISNASVNVLVQNCKIYNDASCDISADGQLLAAFIPSSQRGFPDEGILAVYSLA  927
tr|A0A2I3TB21|A0A2I3TB21_PANTR      TKFDLPEISNASVNVLVQNCKIYNDASCDISADGQLLAAFIPSSQRGFPDEGILAVYSLA  927
sp|A2AH22|AMRA1_MOUSE               TKFDLPEISNASVNVLVQNCKIYNDASCDISADGQLLAAFIPSSQRGFPDEGILAVYSLA  928
tr|F1LNQ2|F1LNQ2_RAT                TKFDLPEISNASVNVLVQNCKIYNDASCDISADGQLLAAFIPSSQRGFPDEGILAVYSLA  928
                                    **:*********:**** ******************.***********************

sp|E7FAG6|AMR1A_DANRE               PHNLGEMLYSKRFGPNAISVSLSPMGRYVMVGLASRRILLHQISDHMVAQVFRLQQPHAG  930
tr|F6YH53|F6YH53_XENTR              PHNLGEILFTKRFGPNAISVSLSPMGRYVMVGLASRRILLHPSTEHMVAQVFRLQKPHGG  999
tr|F1MPW0|F1MPW0_BOVIN              PHNLGEMLYTKRFGPNAISVSLSPMGRYVMVGLASRRILLHPSTEHMVAQVFRLQQAHGG  958
sp|Q9C0C7|AMRA1_HUMAN               PHNLGEMLYTKRFGPNAISVSLSPMGRYVMVGLASRRILLHPSTEHMVAQVFRLQQAHGG  987
tr|A0A2I3TB21|A0A2I3TB21_PANTR      PHNLGEMLYTKRFGPNAISVSLSPMGRYVMVGLASRRILLHPSTEHMVAQVFRLQQAHGG  987
sp|A2AH22|AMRA1_MOUSE               PHNLGEMLYTKRFGPNAISVSLSPMGRYVMVGLASRRILLHPSTEHMVAQVFRLQQAHGG  988
tr|F1LNQ2|F1LNQ2_RAT                PHNLGEMLYTKRFGPNAISVSLSPMGRYVMVGLASRRILLHPSTEHMVAQVFRLQQAHGG  988
                                    ******:*::*******************************  ::**********: *.*

sp|E7FAG6|AMR1A_DANRE               ETSMRRVFDVVYPMAPDQRRHVSINSARWLPDPGLGLAYGTNKGDLVICRPVDVHSDGSS  990
tr|F6YH53|F6YH53_XENTR              ETSMRRVFNVLYPMPADQRRHVSINSARWLPEPGLGLAYGTNKGDLVICRPEAFDNVCDQ  1059
```

```
tr|F1MPW0|F1MPW0_BOVIN              ETSMRRVFNVLYPMPADQRRHVSINSARWLPEPGLGLAYGTNKGDLVICRPEASNSGVEY  1018
sp|Q9C0C7|AMRA1_HUMAN               ETSMRRVFNVLYPMPADQRRHVSINSARWLPEPGLGLAYGTNKGDLVICRPEALNSGVEY  1047
tr|A0A2I3TB21|A0A2I3TB21_PANTR      ETSMRRVFNVLYPMPADQRRHVSINSARWLPEPGLGLAYGTNKGDLVICRPEALNSGVEY  1047
sp|A2AH22|AMRA1_MOUSE               ETSMRRVFNVLYPMPADQRRHVSINSARWLPEPGLGLAYGTNKGDLVICRPEALNSGIEY  1048
tr|F1LNQ2|F1LNQ2_RAT                ETSMRRVFNVLYPMPADQRRHVSINSARWLPEPGLGLAYGTNKGDLVICRPEALNSGIEY  1048
                                    ********:*:***  ***************:*******************   ..  .

sp|E7FAG6|AMR1A_DANRE               TSE-HSERMFTINNGGGVGPSSSRSGDRAGSSRTDRRSRRDIGLMNGVGLQPQPPAASVT  1049
tr|F6YH53|F6YH53_XENTR              FWEQMNEAILQ--------HNTPRSSERPGTSRASWRSDRDMGLINAIGLQPRNPTTSVT  1111
tr|F1MPW0|F1MPW0_BOVIN              YWDQLNETVFTV-------HSSSRSSERPGTSRATWRTDRDMGLMNAIGLQPRNPTTSVT  1071
sp|Q9C0C7|AMRA1_HUMAN               YWDQLNETVFTV-------HSNSRSSERPGTSRATWRTDRDMGLMNAIGLQPRNPATSVT  1100
tr|A0A2I3TB21|A0A2I3TB21_PANTR      YWDQLNETVFTV-------HSNSRSSERPGTSRATWRTDRDMGLMNAIGLQPRNPATSVT  1100
sp|A2AH22|AMRA1_MOUSE               YWDQLSETVFTV-------HSSSRSSERPGTSRATWRTDRDMGLMNAIGLQPRNPTTSVT  1101
tr|F1LNQ2|F1LNQ2_RAT                YWDQLNETVFTV-------HSSSRSSERPGTSRATWRTDRDMGLMNAIGLQPRNPTTSVT  1101
                                      :  .* ::          .. **.:* *:**:  *: **:**:*.:****: *::***

sp|E7FAG6|AMR1A_DANRE               SQGTQTQNQRLQHAETQTDRDLPDDPQQPSTSQG-SQVTDATESLDFETLPEDSGSEVVP  1108
tr|F6YH53|F6YH53_XENTR              SQGTQTPAPQLQNAETQTEREVPESRPAPPTAEAGPSGTAETPSSSSEGA----------  1161
tr|F1MPW0|F1MPW0_BOVIN              SQGTQTLALQLQNAETQTEREIQEPGVAA----SGP------------------------  1103
sp|Q9C0C7|AMRA1_HUMAN               SQGTQTLALQLQNAETQTEREVPEPGTAA----SGP------------------------  1132
tr|A0A2I3TB21|A0A2I3TB21_PANTR      SQGTQTLALQLQNAETQTEREVPEPGTAA----SGP------------------------  1132
sp|A2AH22|AMRA1_MOUSE               SQGTQTLALQLQNAETQTEREEEEPGAAS----SGP------------------------  1133
tr|F1LNQ2|F1LNQ2_RAT                SQGTQTLALQLQNAETQTEREEEEPGTAS----SGP------------------------  1133
                                    ******   :**:*****:*:  :         .

sp|E7FAG6|AMR1A_DANRE               ETPPHSRPQEDEGSDPSEPSTDSTGQAEYVSRIRRLMAEGGMTAVVQREQSTTMASMGSF  1168
tr|F6YH53|F6YH53_XENTR              ---PSSGESQEGIPSSSEVPGSGEGQEDALSRIQRLMAEGGMTAVVQREQSTTMASMGGF  1218
tr|F1MPW0|F1MPW0_BOVIN              --------------GEGEGSDYGASGEDALSRIQRLMAEGGMTAVVQREQSTTMASMGGF  1149
sp|Q9C0C7|AMRA1_HUMAN               --------------GEGEGSEYGASGEDALSRIQRLMAEGGMTAVVQREQSTTMASMGGF  1178
tr|A0A2I3TB21|A0A2I3TB21_PANTR      --------------GEGEGSEYGASGEDALSRIQRLMAEGGMTAVVQREQSTTMASMGGF  1178
sp|A2AH22|AMRA1_MOUSE               --------------GEGEGSEYGGSGEDALSRIQRLMAEGGMTAVVQREQSTTMASMGGF  1179
tr|F1LNQ2|F1LNQ2_RAT                --------------GEGEGSEYGGSGEDALSRIQRLMAEGGMTAVVQREQSTTMASMGGF  1179
                                                  . .*    . .  : :***:************************.*

sp|E7FAG6|AMR1A_DANRE               GNNIIVSHRIHRGSQTGADAQNRTRLSPIPGPSSGAPESLAAASYSRVLTNTLGFRGDTA  1228
tr|F6YH53|F6YH53_XENTR              GNNIIVSHRIHRGSQTASDPASRASAPRSPQPSTSRETVAELE--------RLLAPPQTS  1270
tr|F1MPW0|F1MPW0_BOVIN              GNNIIVSHRIHRSSQTGTEPGAA--HAPSPQPSTSRELLPEAG--------QLTERG---  1196
sp|Q9C0C7|AMRA1_HUMAN               GNNIIVSHRIHRSSQTGTEPGAA--HTSSPQPSTSRGLLPEAG--------QLAERG---  1225
tr|A0A2I3TB21|A0A2I3TB21_PANTR      GNNIIVSHRIHRSSQTGTEPGAA--HTSSPQPSTSRGLLPEAG--------QLAERG---  1225
sp|A2AH22|AMRA1_MOUSE               GNNIIVSHRIHRSSQTGTESGAA--RTSSPQPSTSRGLPSEPG--------QLAERA---  1226
tr|F1LNQ2|F1LNQ2_RAT                GNNIIVSHRIHRSSQTGTESGAA--RTSSPQPSTSRGLLSEPG--------QLAERG---  1226
                                    ************.***.::          * **:.                 *

sp|E7FAG6|AMR1A_DANRE               QGIDLTEQERLHTSFFTPEFSPLFSSAVDATGPSSSIGADSVLEGEDFHDFASLPPSLLS  1288
tr|F6YH53|F6YH53_XENTR              Q-------------LLLPETQLLTNNNNDGEVN-----VQPLSSG---------------  1297
tr|F1MPW0|F1MPW0_BOVIN              ---------------LSPRTASWEQPATPGREP--------ALPSSS----PAPPPAHLP  1229
sp|Q9C0C7|AMRA1_HUMAN               ---------------LSPRTASWDQPGTPGREP-----TQPTLPSSS----PVPIPVSLP  1261
tr|A0A2I3TB21|A0A2I3TB21_PANTR      ---------------LSPRTASWDQPGTPGREP-----TQPTLPSSS----PVPIPVSLP  1261
sp|A2AH22|AMRA1_MOUSE               ---------------LSPRTASWDQPSTSGREL-----PQPALSSSS----PVPIPVPLA  1262
tr|F1LNQ2|F1LNQ2_RAT                ---------------LSPRTASWDQPSTSGREL-----PQPALSSSS----PVPIPVPLA  1262
                                                   : *.     .    .              .

sp|E7FAG6|AMR1A_DANRE               SS-----PSLSPVNNSNYSNSDSSYLGDEYGR-------       1315
tr|F6YH53|F6YH53_XENTR              ----------------TFPGVE---R-------------       1304
tr|F1MPW0|F1MPW0_BOVIN              STEGPTPPRCDLTNSNHLPDSGGSGRGEAAGPSGEPRDR       1268
sp|Q9C0C7|AMRA1_HUMAN               SAEGPT-LHCELTNNNHLLDG-GSSRGDAAGPRGEPRNR       1298
tr|A0A2I3TB21|A0A2I3TB21_PANTR      SAEGPT-LHCDLTNNNHLLDG-GSSRGDAAGPRGEPRNR       1298
sp|A2AH22|AMRA1_MOUSE               SNEGPT-MHCNVTNNSHLPEGDGSNRGEAAGPSGEPQNR       1300
tr|F1LNQ2|F1LNQ2_RAT                SNEGPT-MHCNVTNNSHLPEGDSSNVGEAAGPSGEPRNR       1300
```
